# Drying of virus-containing aerosol particles: Modelling effects of droplet origin and composition

**DOI:** 10.1101/2021.03.25.436946

**Authors:** Michael C. Jarvis

## Abstract

**Background and Purpose:** Virus-containing aerosol droplets emitted by breathing, speech or coughing dry rapidly to equilibrium with ambient relative humidity (RH), increasing in solute concentration with effects on virus survival and decreasing in diameter with effects on sedimentation and respiratory uptake. The aim of this paper is to model the effect of ionic and macromolecular solutes on droplet drying and solute concentration.

**Methods:** Deliquescence-efflorescence concepts and Kohler theory were used to simulate the evolution of solute concentrations and water activity in respiratory droplets, starting from efflorescence data on mixed NaCl/KCl aerosols and osmotic pressure data on respiratory macromolecules.

**Results:** In NaCl/KCl solutions supersaturated total salt concentrations were shown to reach 10-15M at the efflorescence RH of 40-55%, depending on the K:Na ratio. Dependence on K:Na ratio implies that the evaporation curves differ between aerosols derived from saliva and from airway surfaces. The direct effect of liquid droplet size through the Kelvin term was shown to be smaller and largely restricted to breath emissions. Modelling the effect of proteins and glycoproteins showed that salts determine drying equilibria down to the efflorescence RH, and macromolecules at lower RH.

**Conclusion:** Differences in drying behaviour are predicted between breathing, speech and coughing emissions and between droplet size fractions within these. High salt concentrations may inactivate some viruses.

## Introduction

Some respiratory viruses can be transmitted in aerosol form, as well as in larger droplets and surface deposits^1^. Aerosols are conventionally defined as droplets or particles less than 5-10 μm in diameter that, according to Stokes’ Law, remain suspended in still air for minutes or longer^2^. It has been argued^3, 4^ that the size range should be extended to 50-100 μm because turbulence, either in a cough jet^5^ or due to draughts^6^ or convection^7^, keeps larger particles airborne for longer than is predicted by Stokes’ Law.

Despite initial doubts, it is now quite widely accepted that certain viruses including SARS-CoV-2 are transmitted in aerosols, particularly from asymptomatic subjects^8, 9^. Aerosol transmission is most likely in enclosed spaces such as schools, public buildings and transportation^10^. Aerosols can transmit viruses from person to person with transiently inadequate social distancing^4^, but they can also build up, over minutes to hours, throughout the air in an enclosed space so that the risk of infection depends on the duration of emission and exposure, not on distance^11^. In these circumstances the risk depends on Wells-Riley dynamics and is reduced by ventilation with fresh or filtered air and by anything that decreases the viable lifetime of the virus^11^.

The viable lifetime of airborne viruses varies. SARS-CoV-2 remains viable indoors for minutes to hours^12-15^. The rate of inactivation is enhanced by sunlight^16, 17^ and increases with temperature^17, 18^. There is also a significant effect of humidity on SARS-CoV-2 inactivation^12, 17^, but the nature and mechanism of the humidity effect are still poorly understood.

For some viruses inactivation is faster at intermediate levels of relative humidity, around 40%, than at high or very low humidity^19^. It has been suggested that the common cold virus HRV-16, which follows this pattern, loses activity in the supersaturated salt solution when dried from the high RH of emission to just above the precipitation (efflorescence) zone of the salts present, roughly 40% RH^20^. In contrast, for SARS-CoV-2 in aerosol form, the limited data available suggest that the rate of inactivation is low at all RH values tested up to about 50% and increases at higher RH^17^. The rate of inactivation of surface deposits of SARS^21^ and SARS-CoV-2^22, 23^ is lower than aerosols but likewise rises with increasing humidity^24^. This implies that the SARS viruses are more resistant than, for example, HRV-16 to very high salt concentrations, and that whatever its mechanism the inactivation of SARS-CoV-2 is dependent on available water.

During the drying process, the water activity decreases towards equilibrium with the surrounding RH as the concentration of the solutes rises. If the RH is low enough, however, the solutes precipitate and lose their capacity to retain liquid water, at which RH the aerosol droplet dries abruptly^25^. For salts, the *efflorescence RH* at which salts precipitate, as the RH falls, is normally lower than the *deliquescence RH* at which the salt redissolves when the RH is rising^26^. Between the efflorescence RH and the deliquescence RH the solution is supersaturated. In this RH region the droplet is not strictly in equilibrium with the surrounding air: it is common to speak of ‘equilibration’ but this means only the approach towards an unstable quasi-equilibrium, which would change suddenly if precipitation of the salt were nucleated^26^. Water availability, expressed as the water activity *a*_w_, increases with the ambient RH but is also modulated by changing surface energy through the Kelvin effect in small aerosol droplets^25^.

Due to their influence on virus stability, the salt concentrations and water activity in aerosol-sized droplets deserve closer examination. The same is true for surface deposits, particularly because it is not known why viral viability is enhanced in that form^14, 23^. The drying of aerosols when emitted into ambient air has been quite extensively studied^27-29^ and modelled^27, 28, 30^, often with the aim of predicting when droplets, initially large enough to sediment in still air, will shrink enough to remain suspended. For the largest droplets this depends on the kinetics of drying: they fall to the floor before they have time to dry^27, 28^, although the falling and drying times both depend on turbulence^4^. For smaller droplets drying is rapid and it is the equilibrium (strictly, quasi-equilibrium) with the ambient RH that matters^31^.

In the biomedical literature, a droplet that has dried to equilibrium is called a droplet nucleus^1^. Depending on the solids present and the moisture that their hygroscopicity retains, a droplet nucleus may consist of a very concentrated solution, a polycrystalline salt precipitate, a protein gel or amorphous solid, or a combination of these phases; presenting very different environments in which viruses may be inactivated^19^. For example, thermal denaturation depends on mobile water^32^. Also polycrystalline or other solids may refract or absorb daylight, which is known to inactivate SARS-CoV-2^17^.

Much of the published experimentation on droplet drying (cited by^27^) has made use of simplified analogues of the respiratory fluids in which viruses are emitted by breathing, speaking, singing, coughing or sneezing. Sometimes just NaCl solutions have been used. The detailed salt composition of natural aerosols has a profound effect on their drying behaviour^33^. Metzger et al.^25^ give an accessible description of the underlying physics (Kohler theory) as an appendix: their nomenclature is adopted here. In respiratory droplets, proteins and glycoproteins have been recognised to contribute volume and mass to the dried droplet nuclei, but little attention has been paid to other ways in which these polymers might influence the drying process^27, 31^.

Because of the very high salt and polymer concentrations that can be reached when biological aerosols dry at low RH, classical colligative relationships like Raoult’s Law become increasingly unsatisfactory approximations and the relevant physical chemistry becomes necessarily more empirical. In these circumstances direct experimental measurements using real biological fluids may be more informative than theoretical prediction^27^. However, these measurements are technically challenging^27, 34^ and during the pandemic time is short.

This paper describes simulations of the effects of some of the main variables in the composition of virus-containing aerosols on the drying process. Only the drying (efflorescence) direction of RH change is considered, so the focus is on the quasi-equilibrium with RH in the region between the efflorescence RH and the deliquescence RH, where the salt constituents are supersaturated. In view of the uncertainties discussed above, these simulations do not aim at quantitative descriptions of complex, real-life bioaerosols, but may serve to provide some simplifying assumptions and to prioritise variables that deserve experimental investigation.

## Methods

### Modelling mixed NaCl/KCl solutions

Published efflorescence data^26^ in the form of measured droplet area ratios *R*_a_ with the constant droplet area at RH <30% set as unity, were converted to volume ratios *R*_v_ = *R*_a_^3/2^. This approach was justified by a close match (+/- <1%) with the efflorescence and deliquescence RH measured at bulk scale^26^. For RH below the efflorescence point, *R*_v_ = *R*_0_. To obtain absolute salt concentrations the *R*_v_ scale needs to be calibrated. The calibration was attempted in three different ways (1-3) as described below.

1. For RH above the efflorescence point *R*_v_ is equivalent to the growth factor as conventionally defined^25^. *R*_0_ was converted to mass using a solid density ρ_solid_ interpolated between the densities of KCl (1980 Kg m^-3^) and NaCl (2176 Kg m^-3^) according to the molar ratio, with a correction factor of 0.6 to account for void volume in the polycrystalline salt deposits. Solution concentrations were then calculated as equal to *R*_0_.0.6.ρ_solid_ /(*R*_v_ -1).
2. The solute activity coefficient at the highest measured RH was input into Raoult’s Law to calculate the equilibrium salt concentration at that RH, assuming that the effects of NaCl and KCl were additive. The ion activity coefficients were calculated from an exponential function of the form *a*.exp(-*b*.[salt]) + *c*.[salt] + *d* where *a, b, c* and *d* are empirical constants, fitted to published activity coefficient data for solutions of the pure salts^35^.
3. At the deliquescence RH for a pure salt the solution concentration is equal to the known saturation concentration. For the mixed NaCl/KCl solutions for which experimental data^26^ were used, deliquescence is a two-stage process with a eutonic mixture (K mol fraction = 0.3) dissolving first at the lower deliquescence RH, and the excess salt remaining solid until the upper deliquescence RH is reached. For (K mol fraction = 0.2), therefore, the upper deliquescence RH of 74.2%^26^ was considered to correspond to the solubility of NaCl (6.2 molal) and for (K mol fraction = 0.8) the upper deliquescence RH of 79.0%^26^ was considered to correspond to the solubility of KCl (5.5 molal). This approach is similar to that advocated by Metzger et al.^25^.

Calibration methods (1) and (2), which incorporate considerable uncertainties about the shape and density of the dried particles at RH below the efflorescence RH, and the additivity of the salt effects in aerosols, gave higher concentrations than method (3). Method (3) was therefore used in preference.

Method (3) includes a term for the density of the solutions, which was calculated from an empirical function of the form: *b*[Salt]^2^ + *c*[salt] + *d* where the constants *b, c* and *d* were derived by least-squares fitting to published data^36^ interpolated between NaCl and KCl.

The Kelvin term in the predicted drying equilibrium was calculated as a function of droplet diameter using the relationship

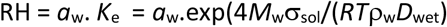

Where *M*_w_ is the molar mass of water, ρ_w_ is its density and σ_sol_ is the surface tension of the solution^25^.

### Modelling water activity in macromolecular solutions

The water activity of protein/glycoprotein solutions without salt was calculated from published osmotic pressure data^37^ using the following form of the Van t’Hoff relationship^38^:

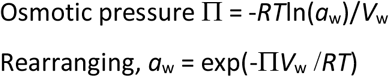

Where RH is expressed as a fraction, *R* is the gas constant, *T* is absolute temperature and *V*_w_ is the molar volume of water.

All simulations were carried out in Microsoft Excel, using the SOLVER function for least-squares fitting. The .xlsx files are available from the author on request.

## Results

### Origins and composition of emitted droplets

Aerosols and larger droplets emitted during breathing^39^, speech^40^, coughing^41^ and other activities^42^ originate by aerodynamic disruption of the mucosal lining^43^ in different zones in the respiratory tract^34, 44^, leading to different droplet size distributions^45^. The principal zones in which droplets are generated are the bronchioli (modal droplet diameter 1-2 μm) the laryngeal region (modal droplet diameter 1-2 μm), and the oral cavity and nasal region (modal droplet diameter >100 μm)^45^. Multiple sites of origin lead to bimodal, trimodal or broad continuous ranges of droplet diameter for each mode of emission^45^. With droplet diameters covering several orders of magnitude, caution is needed in the interpretation of modal figures because they may be derived by several experimental methods with differing size limitations^45^ and because number-weighted and volume-weighted distributions are very different. Volume-weighted distributions are more relevant to viral load^31^.

Droplets emitted in normal breathing are primarily from the bronchial zone and have diameters in the submicron to μm range^45, 46^, while droplets emitted in talking or coughing are derived partly from the laryngeal and oral zones, with a preponderance of larger particles^34, 45^. The viral load of the mucosa in each region^47, 48^ varies with disease progression and between individuals^40^.

The drying of emitted droplets depends on their ionic and polymer composition. It has not been well recognised that the composition of the droplets differs according to their site of origin^27^, and therefore also differs with droplet diameter. A key function of the airway lining throughout the respiratory tract is to sustain hydration^49^ and freedom of motion for the underlying cilia^37^, so defects in hydration lead to disorders such as cystic fibrosis^50^. Equilibrium hydration depends on osmolytes in a very similar way to water retention by emitted droplets^49^, although the RH within the respiratory system is much higher. The cation composition of the airway surface liquid is dominated by sodium, with Na^+^:K^+^ molar ratios typically around 4:1^50-52^: the precision is lower for K^+^ than Na^+^ due to the difficulty of sampling without cellular damage^53^. The principal anion is Cl^-^, with a much smaller amount of HCO_3_^-50, 51^. In health the total salt content is approximately 150 mM^51^, increasing from the lower respiratory tract to the nasal region^50^ and increasing substantially in conditions such as chronic bronchitis^49^. Fluid harvested from human bronchial epithelial (HBE) cell cultures is rather similar in ionic composition to native airway fluids but with lower protein content^52^.

In contrast, saliva has much lower ion concentrations. In the normal (resting) state, the total salt content averages 25 mM and the main cation is K^+^ with a K^+^:Na^+^ ratio of about 3:1^54^. The main anion is Cl^-^. On stimulation, water secretion is driven by an increase in Na^+^ and Cl^-^ concentrations. The Na^+^:K^+^ ratio is therefore variable and can exceed unity^54^. The large difference in salt concentrations between saliva and airway surface fluids means that sputum varies in composition between these extremes^52^. Similarly, emitted droplets are predicted to have an overall K^+^ :Na^+^ ratio that depends on the saliva contribution and is highest for speech^45^, whereas the aerosol droplets emitted by breathing originate mainly from the lining of the lower respiratory tract and are dominated by Na^+^. In emissions of mixed origin, large droplets^45^ are likely to be dominated by K^+^ and small droplets^45^ by Na^+^, leading to differences in their evaporation equilibria.

### Simulated effects of Na^+^:K^+^ ratio

KCl is less soluble than NaCl^26^. Therefore, neglecting the direct (Kelvin) effect of droplet size, large droplets with KCl as the predominant salt would be predicted to reach their efflorescence point at higher RH than small droplets with NaCl as the predominant salt.

However, the behaviour of salt mixtures is not necessarily additive. Li et al.^26^ measured droplet sizes of mixed KCl : NaCl aerosols as they increased with increasing RH and decreased with decreasing RH, using a microscopy technique after impaction.

Figure 1, calculated from the experimental data of Li et al.^26^, shows that in aerosol mixtures of KCl and NaCl, with no other solutes present, the relationship of the minimum water activity at the efflorescence RH to the K:Na ratio is non-linear, with the lowest values reached at about K mol fraction 0.4. Thus each salt tends to keep the other in supersaturated solution until both precipitate together at the efflorescence RH.

**Figure 1.**
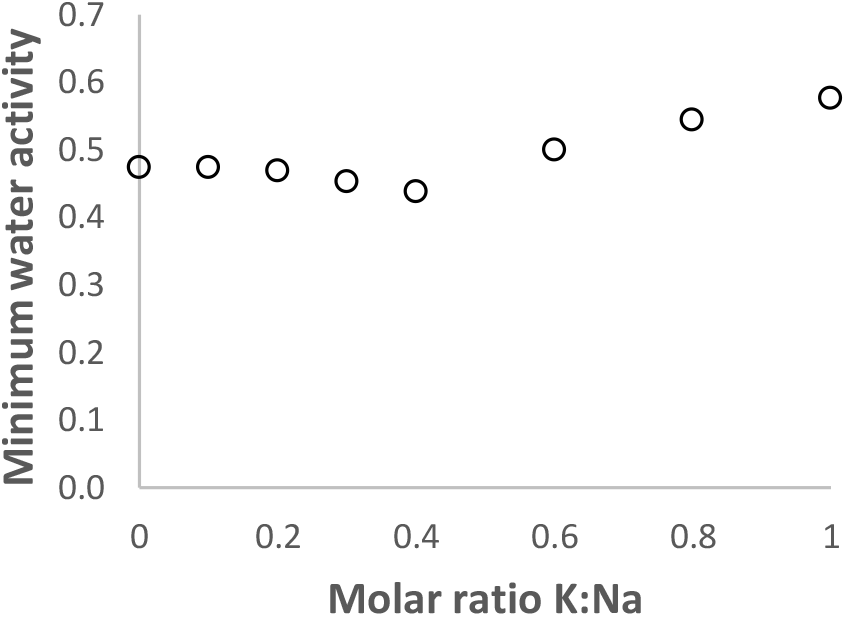
Minimum water activity, at the efflorescence RH, of KCl : NaCl mixtures in aerosol form. Calculated from data in^26^.

Figure 2 shows that In the absence of other solutes, mixtures of NaCl and KCl can reach supersaturated concentrations up to 15 M. Such high concentrations have been suggested to reduce survival of susceptible viruses^20^. Somewhat higher maximal salt concentrations are reached, at lower RH, when the major cation is sodium, mainly because NaCl is more soluble than KCl. When the RH fell below the efflorescence point each droplet contracted abruptly to an irregular solid that remained constant in size down to RH = 5%^26^, from which it was assumed that the liquid phase disappeared at the efflorescence RH.

**Figure 2.**
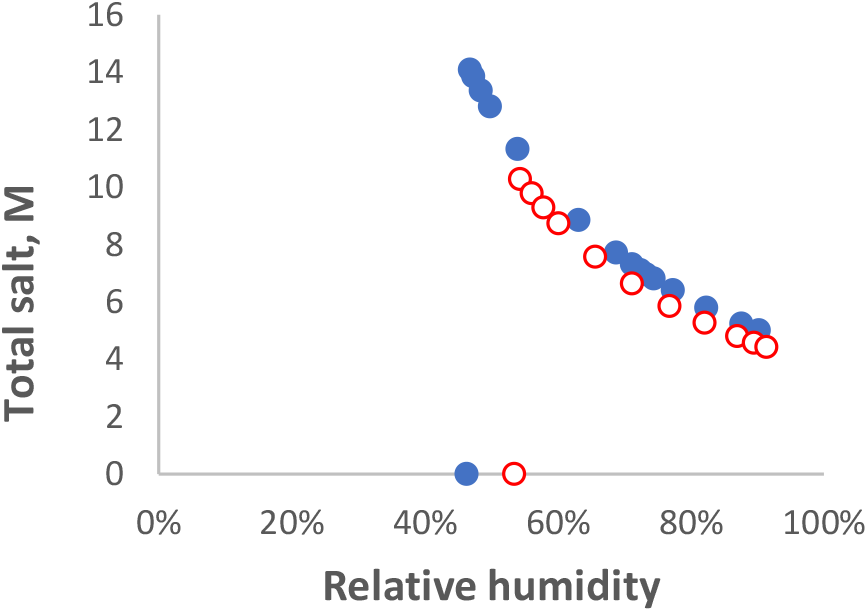
Simulated total salt concentration in aerosols of KCl / NaCl mixtures as a function of decreasing relative humidity, calculated from^26^. Salt concentrations increased as the droplets dried until they reached a maximum at the efflorescence RH. Below the efflorescence RH, with the salts in solid form the droplets dried abruptly.

The mixture with K mol fraction 0.2 is representative of the principal ion content of airway lining fluid^50, 51^ emitted mainly as small (<10 μm) droplets^45^. The mixture with K mol fraction 0.8 is representative of the principal ion content of saliva^54^ emitted mainly as larger (>10 μm) droplets^45^. Ions other than Na^+^ and K^+^ are also present, of course, and their contribution to drying equilibria could be calculated^33^ if comprehensive consensus values for their concentrations were available.

### Droplet diameter: the Kelvin effect

The extent to which droplets dry at any RH is also influenced directly by their size due to the Kelvin effect. The increased surface curvature of small droplets leads to the diameter-dependent Kelvin term *K*_e_ in the expression for their drying equilibrium^25^.

Figure 3 shows that the effect of including the Kelvin term in the simulation is to displace the whole curve to higher RH. The Kelvin effect starts to become ssunstantial only at droplet diameters below about 0.1 μm. In calculating the Kelvin term it is generally assumed that the droplet is wholly liquid and is spherical^25^. When a mixture of irregular solid and liquid phases is present, the local radius at protuberances may be less than calculated and the effect augmented.

**Figure 3.**
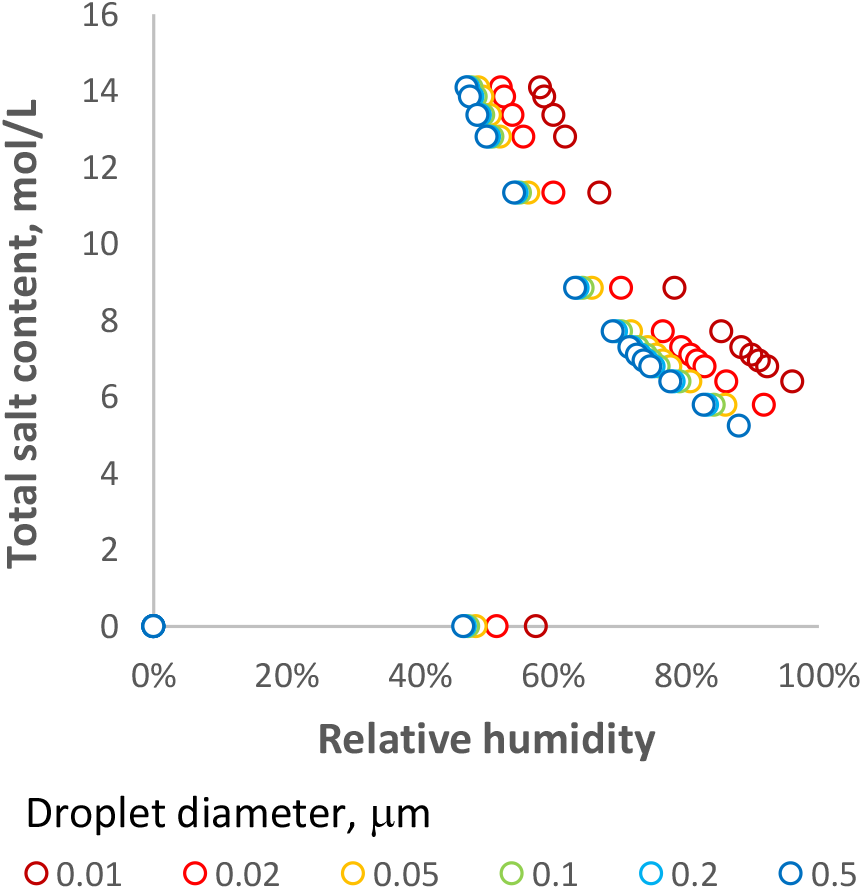
Simulated effect of droplet diameter, through inclusion of the Kelvin term, on the equilibrium salt concentration during drying of aerosol droplets with K mol fraction = 0.2 as in Figure 2.

The magnitude of the Kelvin effect depends also on σ_sol_, the surface tension, which in the simulation above was assumed constant and equivalent to that of water. However, surfactants reduce σ_sol_,^55^ and are present in airway surface liquids^56^, influencing their fragmentation into aerosols^43^. Surfactant proteins^56^ are best known from the lungs but are detectable elsewhere^57^ and are accompanied by deacylated phosphatidyl choline^58^. The effect of surfactants on biogenic droplet drying has not been quantified but it can be assumed that they reduce the magnitude of the Kelvin term, to an uncertain and possibly large extent^55^.

### Macromolecular composition: proteins and mucin glycoproteins

Figure 2 simulates the drying of NaCl/KCl mixtures with no other solutes present. Virus-containing aerosols contain larger amounts of proteins and glycoproteins^52, 54^, which increase in concentration as the droplets dry until they constitute most of the mass of the droplet nucleus. It is often assumed that these macromolecules have no influence on water activity^28^, an assumption that might not be valid when the amount of protein is comparable with the amount of remaining water.

In the intact airway lining, mucin glycoproteins have been stated to play a role in maintaining hydration^49^ and have been studied with that function in mind^52^. Reduced water content and increased viscosity are well-known factors in cystic fibrosis and other pathological conditions^49, 52^. A substantial rheological change attributed to gelation has been observed when the solids content or mucin content of normal airway fluid is doubled^52^, which would occur at RH > 90% during droplet drying. It would then follow that during much of the supersaturation part of the aerosol drying curve the mucin fraction, and perhaps other proteins, are in the gel state. Osmotic relations of polymer gels are difficult to handle, although for simplification it is often assumed that only low-molecular species – free salts and the counterions associated with any charges on the polymer – contribute to lowering water activity^59^. This assumption may not hold at high polymer concentrations or for very flexible polymers that undergo vigorous segmental motion.

In the case of airway fluid, these conceptual problems have been circumvented by direct measurement of the polymer-associated osmotic pressure using a membrane permeable to salts that are not associated with the polymer^37, 49^. An empirical relationship of osmotic pressure to polymeric solids content (proteins plus glycoproteins) of airway mucus, above and below the sol-gel transition, was derived from the measurements of osmotic pressure in ref^37^.

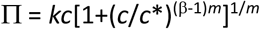

Where *c* is the mucus polymeric solids concentration (Kg/L), *c*\* = 0.081 Kg/L, *m* = 3, β = 2.21 and *k* = 14.4 KPa L/Kg.

Water activity *a*_w_, and hence equilibrium relative humidity, can be connected to osmotic pressure using an appropriate form of the Van t’Hoff relationship^38^:

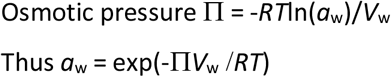

Where *R* is the gas constant, *T* is absolute temperature and *V*_w_ is the molar volume of water.

Figure 4 compares the reduction in water activity due to the salts, calculated (without including the Kelvin term) from the RH data of Li et al.^26^, with the reduction in water activity due to proteins and glycoproteins, calculated from the osmotic pressure data of Button et al^37^ when solutions with ionic and protein/glycoprotein content representative of airway fluid are dried as far as equilibrium with 71% RH.

**Figure 4.**
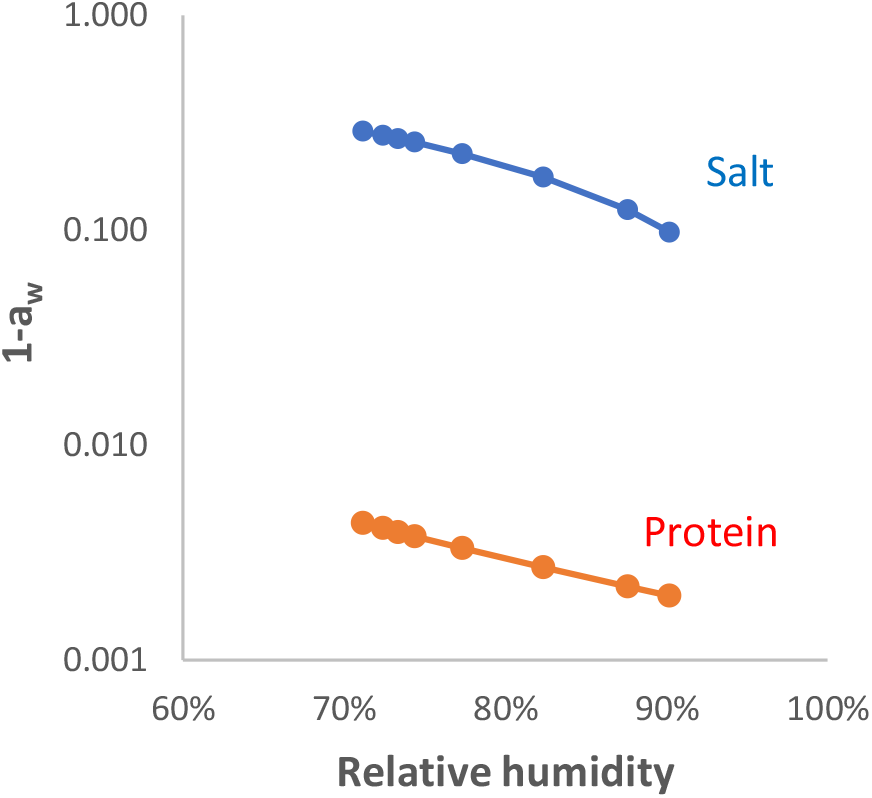
Simulated reduction in water activity a_w_, i.e. (1-*a*_w_), during drying of airway surface liquid by the influence of salt, calculated from RH data for a NaCl/KCl mixture with initial total salt concentration 150 mM and K^+^:Na^+^ molar ratio 0.2 (see Figure 2); and by the influence of the macromolecular (protein and glycoprotein) fraction, calculated from the osmotic pressure measurements of ref^37^. Note that the (1-*a*_w_) scale is logarithmic.

Drying to this extent requires extrapolation of the osmotic pressure data somewhat beyond the measured protein concentration range^37^. Within the concentration range shown, the contribution of the protein and glycoprotein fraction to the reduction in water activity was about two orders of magnitude lower than the contribution of the salt mixture, and so would for many purposes be negligible. Extrapolation further into the concentration range where the mucin components gel would be unsafe, bearing in mind the complexity of the osmotic properties of biomacromolecule gels.

To explore what might happen at lower RH, Figure 5 shows the remaining moisture relative to the mass of protein and glycoprotein (approximated as total solids – (NaCl + KCl)) for compositions representative of airway fluid^52^ (0.2 mol fraction K, 40 g/L total solids) and saliva^54^ (0.8 mol fraction K, 5 g/L total solids). The extent of drying was calculated from the salt content only, neglecting the hygroscopic effect of the polymer fraction. The simulated moisture curves for solutions representing airway fluid and saliva remained above 25% until the efflorescence point was reached. In general proteins at less than 20% moisture form hydrated solids, with the water in adsorbed form retained quite strongly to low RH and no separate liquid phase^60^. This would be expected to happen below the efflorescence point.

**Figure 5.**
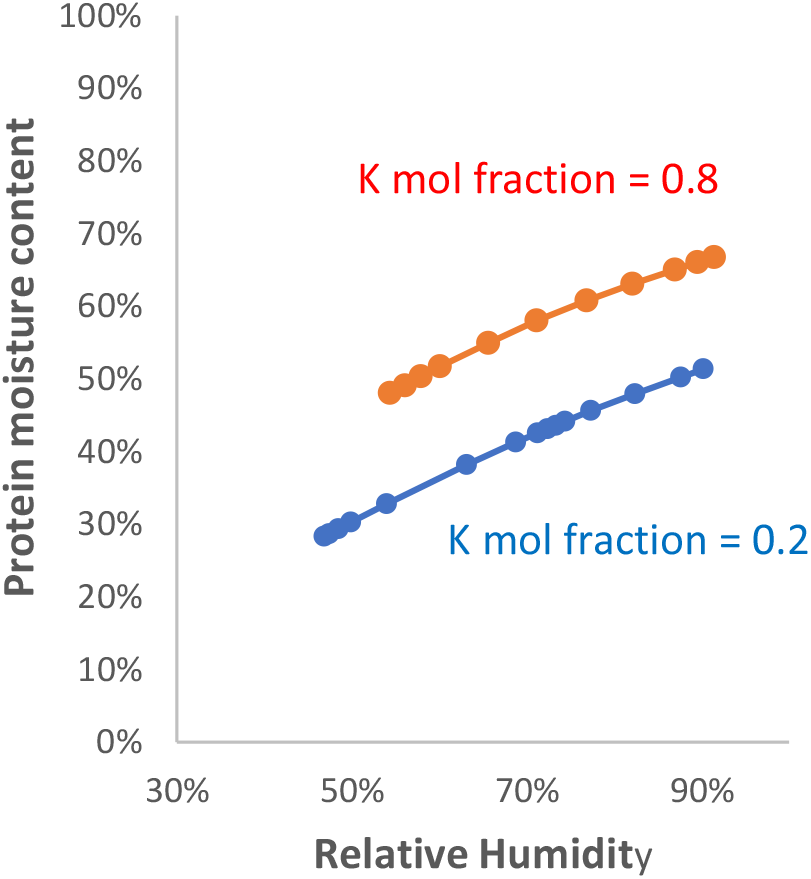
Residual moisture content of the protein and glycoprotein fraction for solutions simulating airway surface liquid (Initial solids 40 g/L; Initial salt 0.15 mol/L; K mol fraction = 0.2) and saliva (Initial solids 5 g/L; Initial salt 0.027 mol/L; K mol fraction = 0.8) during drying.

## Discussion

Supersaturation of salts in aerosol particles means that simple salt mixtures can potentially reach very high concentrations in a rather narrow mid-range band of RH values just above the efflorescence point: below the efflorescence RH the salts precipitate and no longer retain water (Figure 2). It should be noted than anything that can nucleate crystallisation will lead to a lower maximal salt concentration, at higher RH than the efflorescence point. Examples of potential nucleating influences are surfaces, precipitated proteins, or solid particles from pollution aerosols with which the virus-containing droplet has fused. The statistical likelihood of nucleation is greater in larger volumes of solution. Real-world aerosol droplets are therefore likely to show less extreme behaviour than simple salt mixtures, but the observation of clear efflorescence-deliquescence hysteresis in aerosols of model airway fluid^20^ confirms that similar principles apply^33^. The resulting high salt concentrations inactivate some viruses^20^, although SARS-CoV-2 appears to be relatively resistant^17^. Whatever the mechanism by which SARS-CoV-2 is inactivated, the inactivation rate is low in dried droplets and increases with RH, i.e. with the activity of liquid water retained by salts^17^.

Inactivation is particularly slow when droplets dry onto metal surfaces^22^, for reasons that are not clear: water activity in surface deposits depends on ambient RH in the same way as in aerosol droplets large enough for the Kelvin effect to be ignored. Fractionation of solutes during surface drying^61, 62^ might lead to salt-free areas where the virus can survive in dried form at higher RH. There is a potential parallel in the viability of viruses sorbed on atmospheric particulates^31^.

The nature of the cations present in virus-containing emitted droplets deserves closer attention, since the ionic composition of saliva^54^ differs from that of airway fluids^52^. The low salt content and high proportion of KCl in saliva droplets means that their maximal salt content is reached at higher RH than for Na-rich airway fluids. Although a saliva-rich cough^45^ and a sneeze^42^ emit droplets with rather similar initial size ranges, their drying behaviours are predicted to differ.

The dependence of drying equilibrium on droplet size through the Kelvin effect becomes noticeable when the liquid droplet diameter is less than 0.1 μm (Figure 3). Diameter distributions on emission by normal breathing include large numbers of droplets <0.1 μm^45^. However, the volume-weighted diameter distributions, which are more relevant to viral load, include only a small fraction of <0.1 μm droplets, even for breathing^31^. Also, the magnitude of the Kelvin effect is reduced to an unknown extent^55^ by the presence of pulmonary and other surfactants. Thus for many purposes it is a reasonable assumption to neglect the Kelvin term in drying simulations. This prediction has practical consequences. Variations in drying equilibria with droplet size are more likely to arise from differences in site of origin and composition, rather than from the Kelvin effect. Usefully, experiments on the hygroscopic properties of bulk biological fluids, for example measurements of osmotic pressure^37, 49^, are relevant to the behaviour of aerosols so that in suitable cases, it may be possible to access usable experimental data without the technical challenges inherent in aerosol generation and measurement.

However it may be inadvisable to neglect the Kelvin effect for partially dried particles or surface deposits where polycrystalline salts are present, because liquid films may have small local radii overlying crystal vertices and negative (inward) radii in interstices, leading to variation in local water activity.

In the drying range above the efflorescence RH, the hygroscopic effect of proteins and of glycoproteins such as mucins is predicted to be negligible in comparison with the effect of the salts present (Figure 4). Below the efflorescence RH the macromolecular fraction is likely to be essentially an amorphous solid binding the remaining water quite strongly (Figure 5). In the 40% -80% solids range the mucin fraction at least is likely to form a gel, within which water relations are difficult to predict^59^. Other, salted-out proteins may be interspersed with the mucin glycoproteins and such protein aggregates may act as nucleation points for salt precipitation above the efflorescence RH. In this region of complex physical chemistry, experiments on real biological fluids may be a better guide to the behaviour of the system than available theory.

It is concluded that the drying equilibria of aerosol and larger droplets containing infectious viruses are determined principally by the salt composition of the droplets. The salts present in saliva are K^+^-dominated, whereas the more concentrated salts present in airway surface liquids are Na^+^-dominated and, in the absence of other solutes, precipitate (effloresce) at lower RH. These differences mean that droplets emitted by breathing, speech, coughing and sneezing differ in drying behaviour according to their site of origin and that in emissions with multiple sites of origin, drying behaviour differs between small and large droplets.

## Availability of data

Excel spreadsheets deriving the numerical data presented are available on request from the author

## Availability of code

see above.

